# dRNA-seq implicates sulfide as master regulator of S(0) metabolism in *Chlorobaculum tepidum* and other green sulfur bacteria

**DOI:** 10.1101/172254

**Authors:** Jacob M. Hilzinger, Vidhyavathi Raman, Kevin E. Shuman, Brian J. Eddie, Thomas E. Hanson

## Abstract

The green sulfur bacteria (*Chlorobiaceae*) are anaerobes that use electrons from reduced sulfur compounds (sulfide, S(0), and thiosulfate) as electron donors for photoautotrophic growth. *Chlorobaculum tepidum*, the model system for the *Chlorobiaceae*, both produces and consumes extracellular S(0) globules depending on the availability of sulfide in the environment. These physiological changes imply significant changes in gene regulation, which has been observed when sulfide is added to *Cba. tepidum* growing on thiosulfate. However, the underlying mechanisms driving these gene expression changes, i.e. specific regulators and promoter elements involved, have not yet been defined. Here, differential RNA-seq (dRNA-seq) was used to globally identify transcript start sites (TSS) that were present during growth on sulfide, biogenic S(0), and thiosulfate as sole electron donors. TSS positions were used in combination with RNA-seq data from cultures growing on these same electron donors to identify both basal promoter elements and motifs associated with electron donor dependent transcriptional regulation. These motifs were conserved across homologous *Chlorobiaceae* promoters. Two lines of evidence suggest that sulfide mediated repression is the dominant regulatory mode in *Cba. tepidum*. First, motifs associated with genes regulated by sulfide overlap key basal promoter elements. Second, deletion of the gene CT1277, encoding a putative regulatory protein, leads to constitutive over-expression of the sulfide:quinone oxidoreductase CT1087 in the absence of sulfide. The results suggest that sulfide is the master regulator of sulfur metabolism in *Cba. tepidum* and the *Chlorobiaceae*. Finally, the identification of basal promoter elements with differing strengths will further the development of synthetic biology in *Cba. tepidum* and perhaps other *Chlorobiaceae*.

**Importance:** Elemental sulfur is a key intermediate in biogeochemical sulfur cycling. The photoautotrophic green sulfur bacterium *Chlorobaculum tepidum* both produces and consumes elemental sulfur depending on the availability of sulfide in the environment. Our results reveal transcriptional dynamics of *Chlorobaculum tepidum* on elemental sulfur, and increase our understanding of the mechanisms of transcriptional regulation governing growth on different reduced sulfur compounds. This study identifies new genes and sequence motifs that likely play significant roles in the production and consumption of elemental sulfur. Beyond this focused impact, this study paves the way for the development of synthetic biology in *Chlorobaculum tepidum* and other *Chlorobiaceae* by providing a comprehensive identification of promoter elements to control gene expression, a key element of strain engineering.

## Introduction

The green sulfur bacteria (*Chlorobiaceae*) are a family of anaerobic photoautotrophic sulfur oxidizers. *Chlorobaculum tepidum*, the model system for this family, both produces and consumes extracellular S(0) globules depending on the availability of sulfide in the environment (1). Recently, Hanson *et al.* showed that biogenic S(0) globules could serve as the sole photosynthetic electron donor for growth of *Cba. tepidum*, and that cell-S(0) contact was required for that growth (2). Expanding upon this work, Marnocha *et al.* showed that cell-S(0) contact was dynamic, with a large populations of unattached *Cba. tepidum* cells growing at similar rates to attached cells (3). As polysulfides were detected in supernatants from cultures producing and consuming S(0), Marnocha *et al.* proposed a model whereby polysulfides act as soluble intermediates in the formation and degradation of S(0) globules, and could feed unattached cells (3). These two studies have significantly increased our understanding of how *Cba. tepidum* interacts with S(0). However many questions remain, particularly how S(0) metabolism is regulated, and what proteins play a role in cell-S(0) attachment.

The analysis of the *Cba. tepidum* genome noted that it encoded few recognizable transcriptional regulators, leading the authors to conclude that *Cba. tepidum* employs little transcriptional regulation (4). However, Eddie and Hanson observed a complex transcriptional response following the addition of sulfide to *Cba. tepidum* growing on thiosulfate (5). The transcript abundance of a two-gene operon (CT1276-CT1277) was significantly increased in the presence of sulfide, with CT1277 belonging to the helix-turn-helix xenobiotic response element family-like protein superfamily, indicating that transcriptional regulators with limited functional annotation may contribute to gene expression regulation in *Cba. tepidum* (5). While this study identified numerous transcriptionally regulated genes for the transition from thiosulfate to sulfide as a primary electron donor, the *Cba. tepidum* transcriptome during growth on S(0) as a sole electron donor has not been documented.

Here, RNA-seq and differential RNA-seq (dRNA-seq) were used to characterize the transcriptome of *Cba. tepidum* growing on biogenic S(0) and to globally identify transcript start sites (TSS) active during growth on sulfide, biogenic S(0), and thiosulfate as sole electron donors, respectively. RNA-seq data suggest that the most dynamic changes in transcript abundance occur in response to sulfide, and that the majority of genes differentially expressed in response to sulfide are downregulated. TSS positions were used to identify putative promoter elements, and, in combination with RNA-seq data, were used to identify DNA motifs that were conserved across homologous *Chlorobiaceae* promoters. The position of the discovered motifs relative to TSS and basal promoter elements suggests that repression is the dominant mode of transcriptional regulation for genes involved in sulfur metabolism in agreement with the RNA-seq data. In support of this observation, deletion of CT1277 from the *Cba. tepidum* genome led to overexpression of the sulfide:quinone oxidoreductase (SQR) CT1087 during growth on thiosulfate, suggesting that the CT1277 gene product acts as a repressor.

## Results

### The transcriptome of *Cba. tepidum* grown with S(0) as the sole electron donor

RNA-seq analysis of transcriptomes in *Cba. tepidum* cultures growing on thiosulfate and biogenic S(0) revealed similar expression profiles for growth on both substrates. 28,183,709 reads were uniquely aligned to the genome for the thiosulfate library, while the S(0) library had 37,350,690 uniquely aligned reads. Despite the tight correlation of expression between thiosulfate and S(0), 120 genes were differentially expressed, with 55 genes being downregulated on S(0) relative to thiosulfate, and 65 genes being upregulated on S(0) (Table S1). Comparing the S(0) transcriptome with the previously published sulfide transcriptome (5) identified 106 differentially expressed genes, with 35 genes upregulated on sulfide relative to S(0), and 71 genes being downregulated on sulfide (Table S1).

The magnitude of transcript abundance changes was examined in detail across the substrates. Log_2_ fold-change values for genes with increased expression on sulfide relative to thiosulfate and S(0) (15.2 and 15.7, respectively) were both much larger than that of S(0) relative to thiosulfate (5.74). Similarly, the fold-change values for genes with decreased expression on sulfide relative to thiosulfate and S(0) were considerably lower than that of S(0) relative to thiosulfate (-9.31 and -9.85 versus -2.30). This suggests that the most dynamic changes in transcript abundance occur in response to sulfide. These changes are clearly observed in genes encoding key components of *Cba. tepidum*’s sulfur oxidation machinery. For example, the Sox and Dsr genes were strongly downregulated on sulfide (5), but were not differentially expressed between growth on S(0) and thiosulfate (Table S1).

### Global identification of TSSs and basal promoter elements in *Cba. tepidum*

The dRNA-seq protocol used in this study produced 6.1-17 million reads per replicate (Table S2) with 2.0-6.7 million reads uniquely aligned to the genome for each replicate. Analysis of these data identified a total of 3426 putative TSS across growth on three electron donors in *Cba. tepidum*: 1086 primary TSS (pTSS), 393 secondary TSS (sTSS), 583 antisense TSS (asTSS), 2100 internal TSS (iTSS), and 71 orphan TSS (oTSS) (Fig. 1A; Table S3). Four transcript start sites previously identified by primer-extension analysis (*csmB*, *csmC, csmE*, and *sigA*) were captured in these data (6). Additionally, a TSS was identified that corroborates the *csmI* promoter inferred by Gruber and Bryant (6). Although the absolute number of putative TSS for each condition was similar (2458 for sulfide, 2303 for S(0), and 2477 for thiosulfate), there were condition-specific differences between predicted TSS (Fig. 1B). However, there was little correlation between condition-dependent TSS and changes in transcript abundance. A total of 1417 of the predicted TSS were found in all three conditions: 718 pTSS, 102 sTSS, 613 iTSS, 338 asTSS, and 33 oTSS.

**Figure 1.**
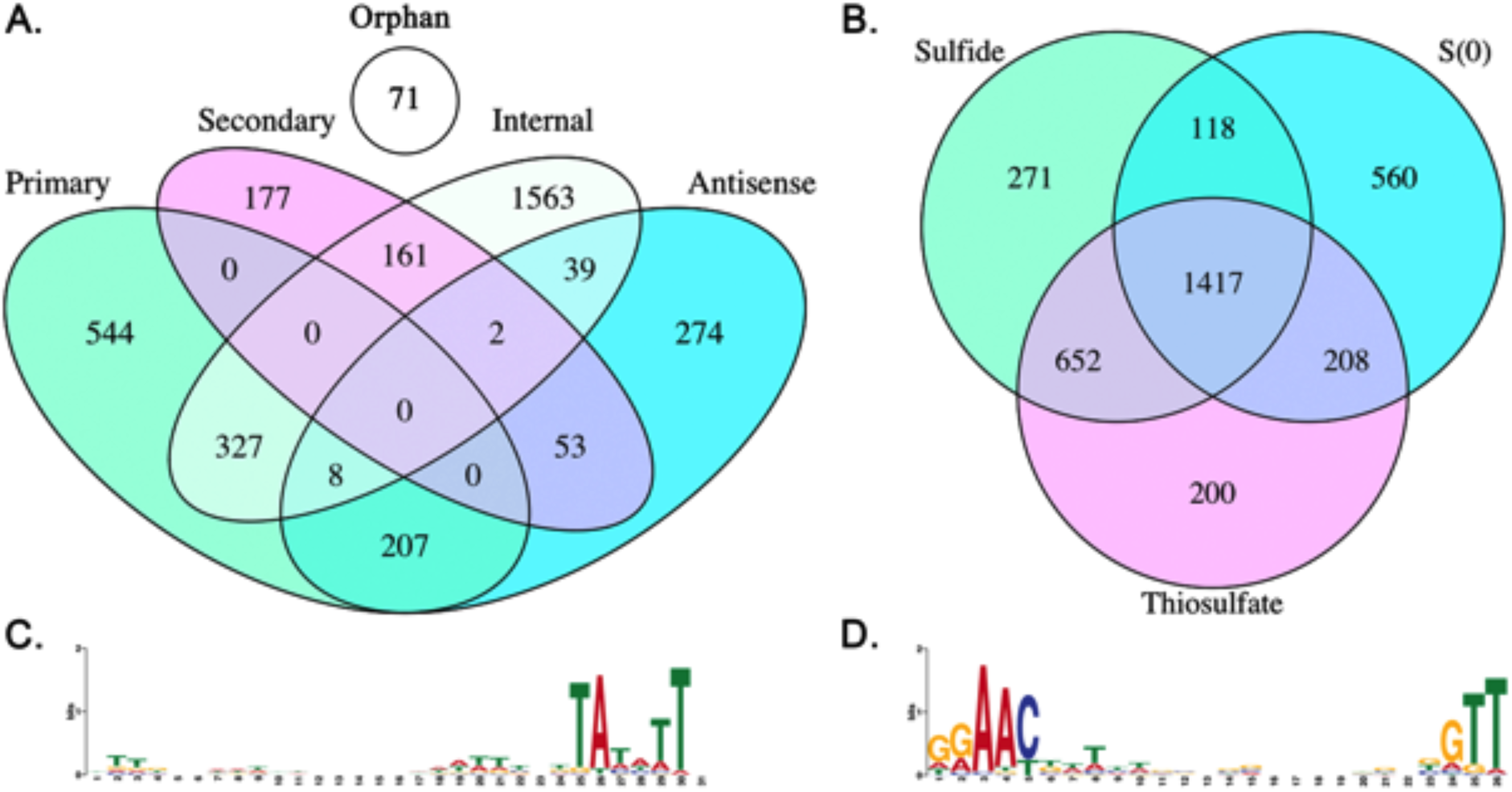
Venn diagrams for TSS data organized by TSS classification (**A**), or by growth condition (**B**). Core promoter motifs identified from analyzing 50 bp upstream of all TSS: motif RpoD motif (**C**) and an ECF factor-like (**D**).

Two motifs were identified when 50 bp upstream of all TSS were analyzed by the MEME software package (Fig. 1C,D; 7). The most abundant motif (Fig. 1C) closely resembles consensus RpoD binding sequences, with the highest similarity to those of the *Bacteroidetes*, specifically *Flavobacterium* spp. (TTG-N_17-23_-TANNTTTG; 8). This would be expected as RpoD protein sequences from *Bacteroidetes* and *Cba. tepidum* clade together, and away from other RpoD proteins (9). However, the -7 consensus sequence, TA[ATC][ATC][AT]T, is different than those of other published RpoD consensus binding sites (8), making the *Cba. tepidum* RpoD consensus binding sequence unique to published consensus sequences to date. The RpoD motif was associated with 1227 TSS (36% of all TSS), and 64% of pTSS. Together, this suggests that the primary sigma factor of *Cba. tepidum* is RpoD encoded by CT1551 (*sigA*).

The second motif (Fig. 1D) resembles σ^70^ ECF subfamily binding sequences, with the distinct AAC motif in the -35 consensus region (10). This motif closely resembles the promoter consensus sequences of σ^E^ (GG[AG][AC]C-N_18_-[CG]GTTg) and σ^H^ ([CG]GGAAc-N_17-_[CG]GTT[CG]) from *Mycobacterium tuberculosis* (11, 12), and that of σ^R^ (GGAAT-N_18_-GTT) from *Streptomyces coelicolor (13)*. As the *Cba. tepidum* genome encodes three σ^70^ ECF factors (CT0278, CT0502, CT0648), this motif may represent a consensus motif that all three ECF factors bind to, or it may represent the consensus sequence of a number of highly similar, yet distinct, motifs that bind each ECF factor with different affinities. For example, σ^X^ and σ^W^ promoter sequences of *Bacillus subtilis* are highly similar, with some promoters binding both proteins, while others bind one, but not the other (14). Thus, this motif likely binds one or more ECF factors in *Cba. tepidum*. This ECF-like motif was associated with 182 TSS (5.3% of all TSS), and 7.9% of pTSS.

Aside from RpoD and the three ECF sigma factors, *Cba. tepidum* encodes one additional sigma factor, CT1193 (*rpoN*), a σ^54^ factor that controls expression of nitrogen regulated genes, including the *nif* operon that encodes proteins for N_2_ fixation in diverse bacteria. Phylogenetic footprinting of the *nifH* promoter across the *Chlorobiaceae* identified RpoN and NtrC binding motifs (data not shown; 32, 33). Using the FIMO search tool (15) in the MEME suite with the RpoN motif as a query returned one additional pTSS with a q-value <0.05 preceding gene CT0644, which encodes an HSP20 family protein.

### Putative sulfide operator sequence 1 (PSOS-1) is associated with *sqr* and putative regulatory genes

*Cba. tepidum* displays a robust transcriptional response depending on the reduced sulfur compound utilized for growth. Binning genes by expression pattern, e.g. all genes with increased transcript abundance on sulfide relative to thiosulfate, and then analyzing regions upstream of the associated pTSS did not produce any significant motifs associated with the promoter regions (data not shown). Therefore, we turned to conservation across *Chlorobiaceae* genomes, i.e. phylogenetic footprinting, to identify putative regulatory motifs associated with sulfur regulated genes that are shared between *Cba. tepidum* and other *Chlorobiaceae*.

Phylogenetic footprinting identified an unknown motif (putative sulfide operator sequence 1; PSOS-1) in the promoter of CT1277 (Fig. 2A), a putative transcriptional regulator that was highly upregulated on sulfide compared to thiosulfate (5) and S(0) (Fig. 2). PSOS-1 was found associated with the pTSS for the two bona fide SQRs CT0117 and CT1087 via FIMO searches (15). Phylogenetic footprinting recovered PSOS-1 motifs associated with *sqr* genes that encode sulfide:quinone oxidoreductase across the *Chlorobiaceae*, including all CT1087 homologues (Fig. 2B), and a subset of CT0117 homologues (Fig. 2C). A consensus motif for *Cba. tepidum* PSOS-1 sites was constructed from CT1277, CT0117, and CT1087 sites (Fig. 2E), and used as the seed in a FIMO search against the *Cba. tepidum* genome. In addition to returning the input loci, PSOS-1 was predicted to occur near 156 TSS. The only other TSS with a q-value < 0.05 was a pTSS for CT0742 (Fig. 2D), a putative transmembrane protein in the TauE-like family of anion transporters. In *Neptuniibacter caesariensis*, TauE has been proposed as a sulfite transporter (16). Based on our TSS data, it appears that CT0743 is co-transcribed with CT0742. The CT0743 gene product is a hypothetical protein containing a domain of unknown function.

**Figure 2.**
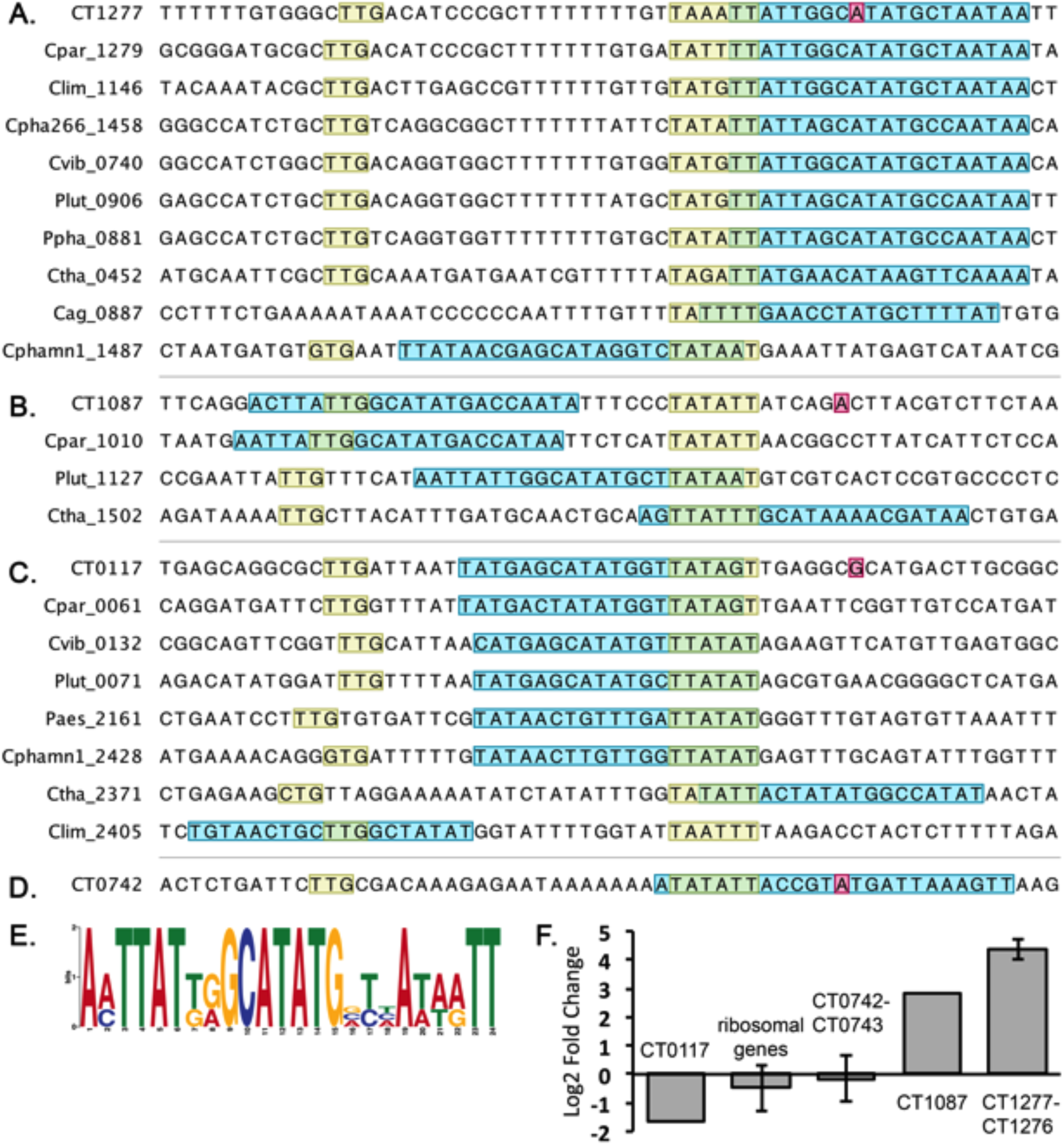
Identification of putative sulfide operator sequence 1 (PSOS-1). Promoter regions for orthologs of CT1277 (**A**) CT1087 (**B**) and CT0117 (**C**) were extracted from all *Chlorobiaceae* genomes. RpoD (yellow) and PSOS-1 (blue) motifs were discovered by promoter analysis. TSS mapped in *Cba. tepidum* are shown in pink. The *Cba. tepidum* consensus motif was searched against the *Cba. tepidum* genome, which returned an additional TSS associated with PSOS-1 (**D**). The PSOS-1 consensus motif for the three *Cba. tepidum* sites (**E**). Log_2_ fold change in transcript abundance on sulfide relative to S(0) of genes associated with PSOS-1 with ribosomal protein genes as a comparator as described in Eddie and Hanson (5) (**F**).

In *Cba. tepidum*, the putative regulator CT1277 appears to be expressed from an RpoD pTSS (Fig. 2A). In the CT1277 promoter, the PSOS-1 motif overlapped the +1 site, and the first two base pairs of the -6 box. The positioning of PSOS-1 relative to the -6 box was conserved for all genomes analyzed except *Chlorobium phaeobacteroides* BS1, where it was found largely in the spacer sequences between the -6 and -33 boxes, with partial overlap of the -6 box (Fig. 2A). In *Chlorobium chlorochromatii*, the PSOS-1 motif was discovered in the promoter of Cag_0887, a hypothetical protein upstream of the CT1277 homologue Cag_0886; Cag_0887-Cag_0885 are likely transcribed as a single unit. The CT1087 and CT0117 PSOS-1 motifs were positioned near the pTSS for each gene (Fig. 2B,C). The positioning of PSOS-1 within homologous CT1087 promoters was variable, with PSOS-1 overlapping the -33 box of two promoters, and the -6 box of the other two. PSOS-1 overlapped the -6 box and extended upstream into the spacer sequence of the six homologous CT0117 promoters. However in the *Chloroherpeton thalassium* promoter, PSOS-1 overlapped the -6 box, and extended downstream, and in the *Chlorobium limicola* promoter, it overlapped the -33 box. Given that the PSOS-1 motif overlaps the RpoD binding motif, the +1 site, or both, suggests that this motif binds a negative repressor, as occlusion of the RpoD binding site, or +1 site, would likely interfere with transcription initiation. The PSOS-1 sequence appears to be unique in that it does not match with any characterized motifs present in the CollecTF, Prodoric, and RegTransBase databases (17).

Expression of the components of the PSOS-1 regulon were variable between growth on sulfide and S(0) (Fig. 2F). CT1277 and CT1087 were both significantly upregulated on sulfide relative to thiosulfate and S(0), while CT0117 was downregulated on both electron donors relative to sulfide (Fig. 2F; 5). CT0742-CT0743 did not change in expression significantly between growth conditions, and displayed a similar expression profile to ribosomal genes.

### PSOS-2 is associated with *sox*, *dsr*, and CT2230 TSS

As many genes related to thiosulfate and S(0) oxidation were found to be downregulated on sulfide relative to S(0) and thiosulfate (Fig. 3; 5), we searched the promoters of genes downregulated on sulfide for putative regulatory motifs. Only one RpoD TSS was observed in the *sox* operon immediately adjacent to *soxJ* (CT1015), supporting the assertion that the *sox* operon is transcribed as a single unit (18). Therefore, sequences upstream of the start codon of the first gene in the *sox* operon across the *Chlorobiaceae* were analyzed for motif discovery. Phylogenetic footprinting identified an unknown motif (putative sulfur operator sequence 2; PSOS-2) in the promoters analyzed (Fig. 3A). The *sox* PSOS-2 motif was searched against the *Cba. tepidum* genome. The top two positions occurring near TSS, excluding the pTSS for *soxJ*, were pTSS for *dsrC-1* (CT0851) and CT2231. Phylogenetic footprinting of *dsrC-1*, *dsrC-*2 (CT2250), and orthologs across the *Chlorobiaceae* returned PSOS-2 sites in all *dsrC* promoters across the *Chlorobiaceae* (Fig. 3B). CT2231, a hypothetical protein with no apparent homologues, appears to be co-transcribed with CT2230, a putative outer membrane protein. As no TSS were found between these two genes, and as both are downregulated on sulfide relative to thiosulfate and S(0) (Fig. 3; 5), sequences upstream of CT2230 homologues across the *Chlorobiaceae* and the CT2231 promoter were analyzed for motif discovery. PSOS-2 sites were found in all promoters (Fig. 3C). A consensus motif for PSOS-2 was constructed from *soxJ*, *dsrC-1*, *dsrC-2*, and CT2231 sites (Fig. 3E). In addition to returning the input loci, PSOS-2 was predicted to occur near 184 additional TSS. The only TSS with a q-value < 0.05 was the pTSS for CT1072 (*dsbE*; Fig. 3D), and an asTSS for CT1231, a transposase annotated as having an internal deletion (data not shown).

**Figure 3.**
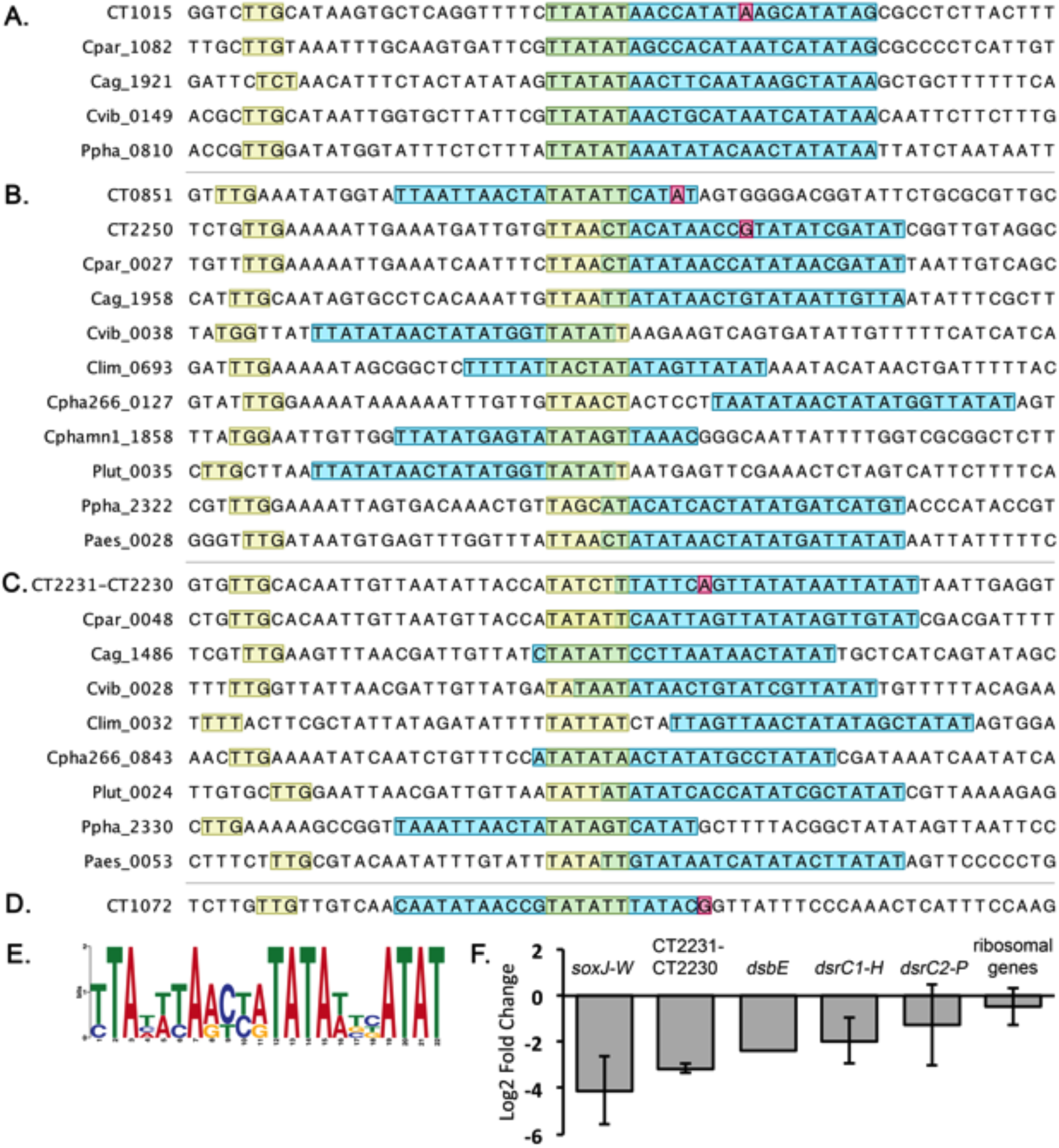
Identification of putative sulfide operator sequence 2 (PSOS-2). Promoter regions for orthologs of the *sox* operon (**A**) *dsrC* (**B**) and CT2230 (**C)** were extracted from all *Chlorobiaceae* genomes. RpoD (yellow) and PSOS-2 (blue) motifs were discovered by promoter analysis. TSS mapped in *Cba. tepidum* are shown in pink. The *Cba. tepidum* consensus motif was searched against the *Cba. tepidum* genome, which returned additional TSS associated with PSOS-2 (**D**). The PSOS-2 consensus motif for the four *Cba. tepidum* sites (**E**). Log_2_ fold change in transcript abundance on sulfide relative to S(0) of genes associated with PSOS-2 with ribosomal protein genes as a comparator as described in Eddie and Hanson (5) (**F**).

In *Cba. tepidum*, PSOS-2 was found to overlap the +1 site of the pTSS of *soxJ*, and the -6 box of the RpoD motif predicted using the bulk TSS data (Fig. 3A). The positioning of PSOS-2 across *sox* promoters of the *Chlorobiaceae* was highly conserved, with all PSOS-2 sites found to overlap the -6 box, and extend downstream. PSOS-2 overlapped both +1 sites of *dsrC-1* and *dsrC-2*, and partially overlapped the -6 box of the RpoD motif (Fig. 3B). The position of PSOS-2 was variable across *dsrC* promoters, yet at least partially overlapped the -6 box in all promoters except that of *Chl. phaeobacteroides*, where it was found 6 bp downstream of the -6 box. PSOS-2 was found to overlap the +1 site of CT2231, and the last bp of the -6 RpoD box (Fig. 3C). Positioning of PSOS-2 was fairly conserved across CT2230 promoters, with all but one site at least partially overlapping the -6 box. The PSOS-2 site in the CT1072 promoter overlapped the -6 box, and part of the RpoD spacer sequence (Fig. 3D). The PSOS-2 sequence appears to be unique in that it does not match with any characterized motifs present in the CollecTF, Prodoric, and RegTransBase databases (data not shown; 17).

The components of the PSOS-2 regulon were downregulated on sulfide relative to S(0) (Fig. 3F). The *sox* operon, CT2231-CT2230, *dsbE, dsrC1A1*, and *dsrC2* are significantly downregulated on sulfide relative to S(0) and thiosulfate (Table S1; 5). The first *dsr* cluster, *dsrCABLEFH* (CT0851-CT0857), appears to be a single transcriptional unit under the control of the *dsrC1* RpoD pTSS. The second *dsr* cluster, *dsrNCABLUEFHTMKJOPVW* (CT2251-CT2238), appears to be broken up into three transcriptional units: *dsrN*, *dsrC2-P,* and *dsrVW*. *dsrN* and *dsrC2-P* appear to be controlled by independent RpoD pTSS, while *dsrVW* appears to be controlled by an ECF factor pTSS (Table S3). This may explain why *dsrVW* is upregulated on sulfide relative to thiosulfate and S(0), while the rest of the cluster is downregulated on sulfide relative to thiosulfate and S(0) (Fig. 3F; Table S1; 5).

### A CRP-like motif (CLM) is associated with *psrABC*, *cydAB* and CT0729

The putative polysulfide oxidoreductase complex, *psrABC* (CT0496-CT0494), is significantly upregulated on sulfide compared to S(0) and thiosulfate (Fig. 4; 5). Phylogenetic footprinting identified two occurrences of a small motif (CRP-like motif; CLM) upstream of the RpoD binding site in the *Cba. tepidum* and *Cba. parvum* promoters (Fig. 4A). The *psrABC* CLM was searched against the *Cba. tepidum* genome for positions that occurred within 200 bp upstream or 50 bp downstream of a TSS. Aside from returning the two *psrABC* sites, FIMO returned positions near the pTSS for CT1818 and CT0729 as the only positions with q-values < 0.05. CT1818-CT1819 encode *cydAB*, a terminal oxidase cytochrome *bd* complex that confers sulfide-resistant O_2_-dependent respiration in *E. coli* (19). CT0729 is a Nudix hydrolase domain protein. A consensus motif was generated from the *psrABC*, *cydAB*, and CT0729 sites, and was used to search against the *Cba. tepidum* genome (Fig. 4C). Significant positions (q-value <0.05) included pTSS for CT1089 and CT1061, and an oTSS at position 503855 (Fig. 4B). CT1089-CT1088 encode for the two subunits of ATP:citrate lyase. CT1061 is a hypothetical protein with a predicted steriodogenic acute regulatory protein-related lipid transfer (START) domain found in polyketide cyclases and dehydrogenases. The sequence downstream of the oTSS was searched for open reading frames, and RNA families (20), but nothing of significance was found.

**Figure 4.**
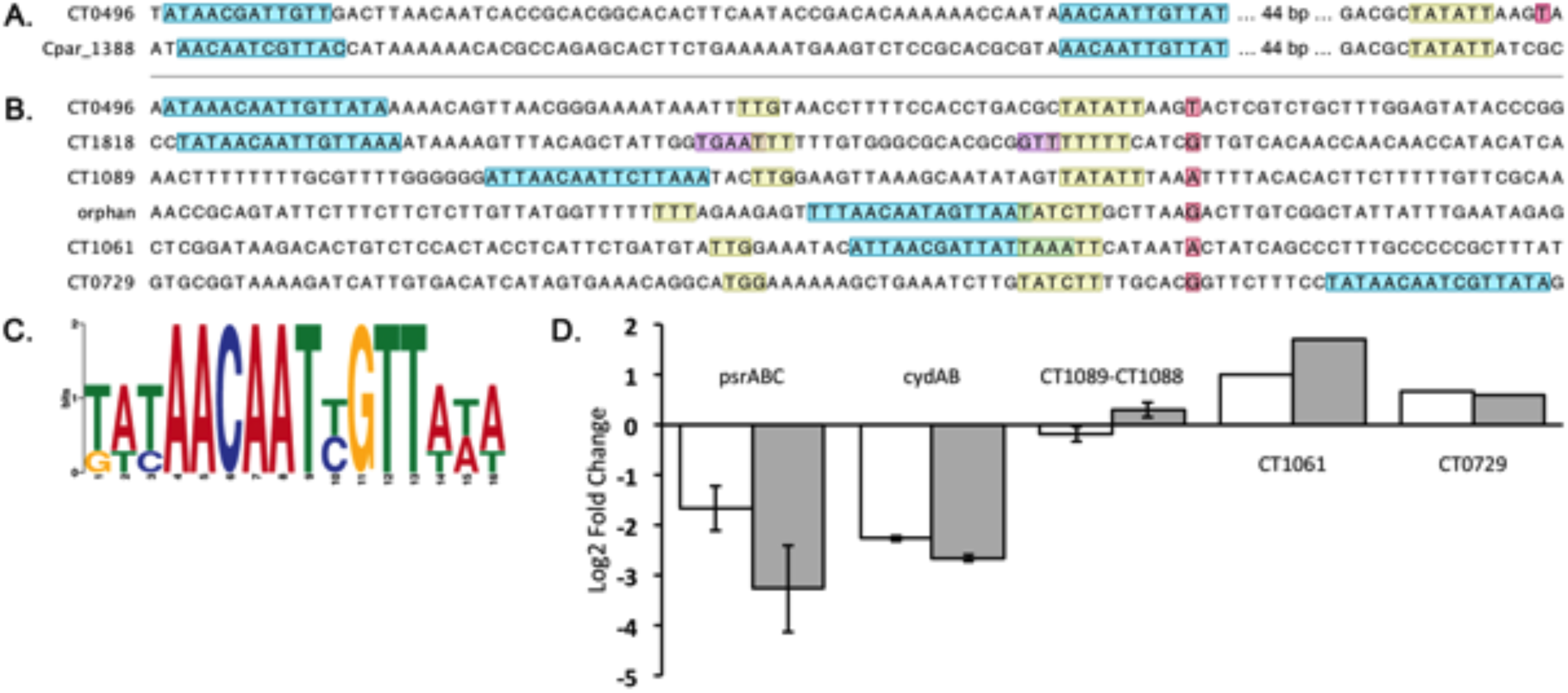
Identification of the CRP-like motif (CLM). Promoter regions for orthologs of the *psrABC* operon were extracted from all *Chlorobiaceae* genomes. RpoD (yellow), ECF sigma factor (purple), and CLM (blue) motifs were discovered by promoter analysis (**A**). TSS mapped in *Cba. tepidum* are shown in pink. The *Cba. tepidum* consensus motif was searched against the *Cba. tepidum* genome, which returned additional TSS associated with CLM (**B**). Consensus motif derived from *Cba. tepidum* CLM sites (**C**). Log_2_ fold change in transcript abundance of genes associated with CLM; S(0) relative to thiosulfate (white) and sulfide relative to S(0) (gray) (**D**).

In *psrABC* promoters of *Cba. tepidum* and *Cba. parvum*, the first CLM site occurred 49 bp upstream of the RpoD -6 box, while the second site occurred 114 bp upstream of the *Cba. tepidum* -6 box, and 113 bp upstream of the *Cba. parvum* -6 box (Fig. 4A). In *Cba. tepidum*, the RpoD -6 box occurs 4 bp upstream of the +1 site. The CLM site for *cydAB* occurred 57 bp upstream of the TSS, in a similar position to the first *psrABC* site that occurred 58 bp upstream of the *psrABC* TSS (Fig. 4B). The CLM site for CT1089 occurred 35 bp upstream of the TSS, and 4 bp upstream of the -33 box, while both the CT1061 and orphan sites overlapped the RpoD spacer sequence and -6 box. The CT0729 CLM site occurred 10 bp downstream of the CT0729 TSS.

The CLM motif resembles the CRP (TGTGA-N_6_-TCACA) and FNR (TTGAT-N_4-_ATCAA) consensus binding motifs of *E. coli* (21, 22) without the spacer sequence. The spacing of the motif sites (Fig. 5A) echoes CRP-activating promoters in *E. coli* with multiple CRP binding sites upstream of the RpoD binding site (21). In support of this, TOMTOM (17) matched the CLM consensus to the PrfA consensus motif from *Listeria monocytogenes*, a CRP-domain transcriptional regulator. The PrfA consensus, WTAACAWWTGTTAA (23), does not contain the N_4-6_ spacer present in *E. coli* CRP/FNR domain binding sites, adding further support that the *Cba. tepidum* CLM may bind a CRP domain protein. *Cba. tepidum* encodes a single CRP/FNR domain protein, CT1719. However, no CRP/FNR domain protein was found in *Cba. parvum*, or any other *Chlorobiaceae* genome examined.

**Figure 5.**
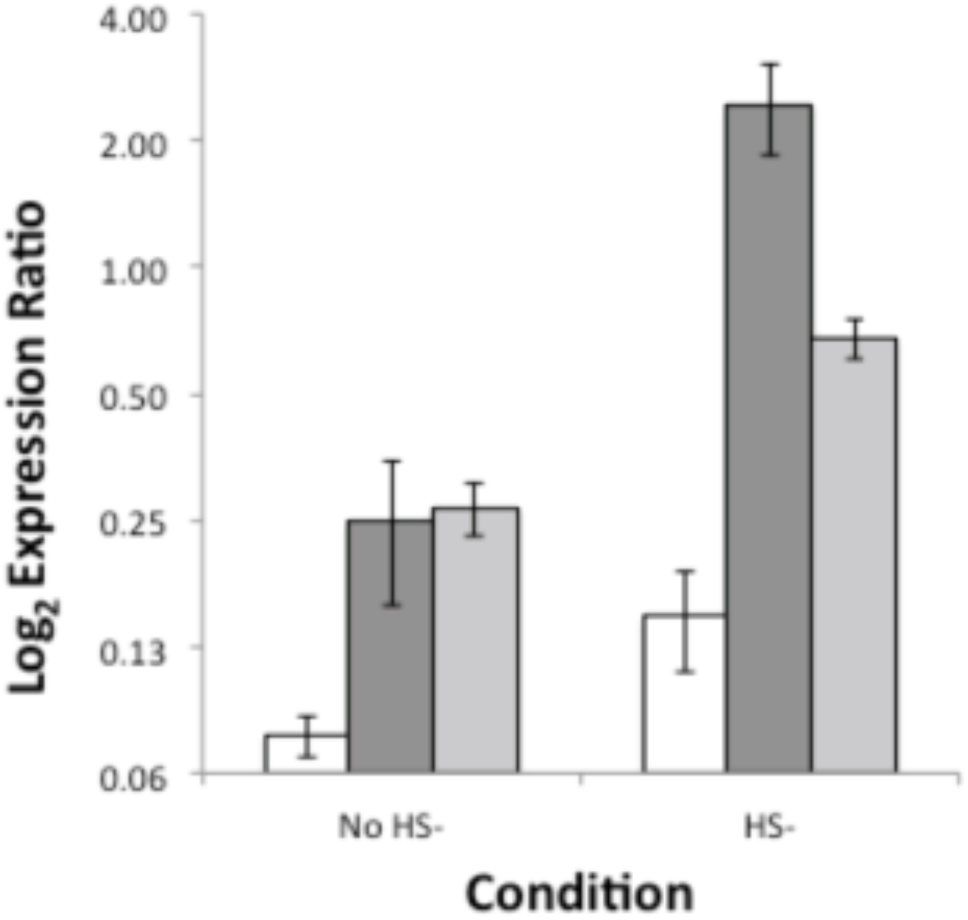
CT1277 is a transcriptional repressor of CT1087. Q-RT-PCR of CT1087 in *Cba. tepidum* wild-type (white), Δ1277.6 (dark gray), and ΔCT1277.11 (light gray) pre-sulfide addition and 40 min post-sulfide addition. Error bars are equal to one standard deviation.

While the components of the CLM regulon have variable expression on S(0) relative to thiosulfate and sulfide, the direction of expression on S(0) relative to sulfide and thiosulfate is similar for each component; *psrABC* and *cydAB* are both significantly downregulated on S(0) relative to thiosulfate and sulfide, while CT1061 and CT0729 are upregulated on S(0) relative to thiosulfate and sulfide (Fig. 4D). CT1089-CT1088 is not differentially expressed between growth conditions.

### CT1277 encodes a transcriptional repressor of CT1087

Previous data showed that the CT1276-CT1277 cassette displayed increased transcript abundance in cells following sulfide addition after growth on thiosulfate (5). Given that CT1277 belongs to the HTH-XRE family of transcriptional regulators, the CT1277 gene was deleted from the *Cba. tepidum* genome to assess its role in sulfide dependent gene regulation. Expression of CT1087 was monitored by q-RTPCR in the wild-type and two independently isolated ΔCT1277 strains (Fig. 5). CT1087 transcript abundance was significantly elevated (3.5-fold, p < 0.03) in both ΔCT1277 strains compared to the wild type during growth on thiosulfate. Transition to growth on sulfide from thiosulfate resulted in a 1.9-fold increase (p = 0.04) in CT1087 transcript abundance in the wild type strain, confirming that its expression is sulfide dependent as previously reported (5, 24). Post-sulfide addition, CT1087 displayed a 16.3-fold (p = 0.003) and 4.6-fold (p < 0.001) increase in transcript abundance relative to the wild type in ΔCT1277.6 and ΔCT1277.11, respectively. Thus, CT1087 expression was much more strongly induced (9.7-fold; p = 0.003) in the absence of CT1277. Increased expression of CT1087 after sulfide addition in the ΔCT1277 strains suggests the presence of a sulfide-dependent activator. Sequences up to 2000 bp upstream of the CT1087, CT1277, and CT0117 promoters were analyzed for motif discovery, but no candidate activator motif was found. Altered expression of CT1087 in the ΔCT1277 strains indicates that CT1277 negatively regulates the transcription of CT1087 in the presence of an activator. No significant growth phenotype was observed for the ΔCT1277 strain compared to the wild-type strain (data not shown).

## Discussion

In this study, we provide the first global transcript abundance data for *Cba. tepidum* during growth on S(0) as the sole electron donor, and use this data to identify genes that were differentially expressed between growth on sulfide and S(0). Many of these genes encode key components of *Cba. tepidum*’s sulfur oxidation machinery. The most dynamic changes in gene expression between growth on different reduced sulfur compounds were in response to sulfide. We also provide the first global transcript start site map for *Cba. tepidum*. dRNA-seq identified 3426 putative TSS across growth on sulfide, thiosulfate, and S(0), of which 1086 were primary TSS. This data also includes 71 orphan TSS that may control transcription of functional elements that were missed during genome annotation, and encompasses the first evidence of widespread antisense transcription in *Cba. tepidum*. Two basal promoter motifs were identified: an RpoD and an ECF sigma factor motif. Three putative regulatory motifs were discovered by phylogenetic footprint analysis of orthologous promoters across the *Chlorobiaceae* for genes that were differentially expressed between growth on reduced sulfur compounds in *Cba. tepidum*. Together, the data presented in this study provides the first predictions for a mechanistic understanding of transcriptional regulation between sulfur-dependent growth states in *Cba. tepidum*, and the *Chlorobiaceae* as a whole.

TSS that are not associated with the RpoD or ECF factor motifs may not bind σ factors, and therefore may not be *bona fide* TSS. In support of this, sequences (+/- 50 bp) surrounding TSS associated with σ factor motifs have significantly higher AT content than those sequences surrounding TSS without σ factor motifs (data not shown). Alternatively, these TSS may be associated with divergent σ factor binding sites that are not represented by the motifs discovered in this study (Fig. 1). Of the 2049 TSS that do not have σ factor motifs, 1733 (84.6%) are iTSS and 319 (15.6%) are pTSS, whereas of the 1377 TSS with σ factor motifs, 367 (26.7%) are iTSS and 767 (55.7%) are pTSS. iTSS are enriched in TSS with no associated σ factor motif, suggesting that most iTSS in this study are not *bona fide* TSS. This is in contrast to a recent study in *E. coli* that found most iTSS were associated with σ factor motifs (27). Indeed, the fraction of iTSS detected in this study, 61% of all TSS, is higher than in other organisms: 37% in *Escherichia coli*, 36% in *Campylobacter jejuni*, and 18% in *Helicobacter pylori* (27, 28, 32). pTSS and sTSS occur at similar percentages across these organisms, while *Cba. tepidum* has fewer asTSS, 17%, than others: 43% in *E. coli*, 48% in *C. jejuni*, and 41% in *H. pylori*. These variations may reflect differences in how more closely related organisms in the *Gamma*- and *Epsilon-proteobacteria* regulate transcription relative to the *Chlorobiaceae*. Future experiments will determine if the high percentage of iTSS in *Cba. tepidum* is characteristic of all *Chlorobiaceae* and if there is any transcriptional activity associated with iTSS associated sequences that lack a recognizable sigma factor motif in *Cba. tepidum*.

A putative operator sequence, PSOS-1, was discovered in the promoters of the two SQRs CT0117 and CT1087, as well as in the promoters of the putative regulatory protein CT1277 and the TauE-domain protein CT0742 (Fig. 2). While CT1087 and CT1277 are highly upregulated on sulfide relative to S(0) (Fig. 2) and thiosulfate (5), CT0117 is downregulated, and CT0742-CT0743 are not differentially expressed. The sulfide versus S(0) expression data corroborates previous studies that have observed downregulation of CT0117, and upregulation of CT1087 and CT1277 in response to high sulfide (5, 24). CT1277 was shown to negatively regulate CT1087 expression (Fig. 5). Therefore, PSOS-1 may represent the CT1277 binding motif, which would suggest that CT1277 is subject to auto-repression. If PSOS-1 binds CT1277, and functions as an operator sequence, then promoters associated with PSOS-1 should be downregulated in response to sulfide; this is the case for CT0117 (Fig. 2). However CT1087 expression increased in response to sulfide in the ΔCT1277 background, suggesting the presence of a sulfide-dependent activator (Fig. 5). As CT1277 is associated with PSOS-1, and shares a similar expression profile to CT1087, then it suggests that the activator acting on CT1087 also activates CT1277. Thus, whatever element activates CT1087 and CT1277 expression should be absent from the CT0117 promoter. This leads to a model whereby sulfide activates the PSOS-1-binding protein (CT1277) and a sulfide-dependent activator (Fig. 6). CT1277 represses transcription of CT0117, CT1087, and CT1277, while the sulfide-dependent activator activates transcription of CT1087 and CT1277, but not that of CT0117. This may be to fine tune expression of CT1277 and CT1087.

**Figure 6.**
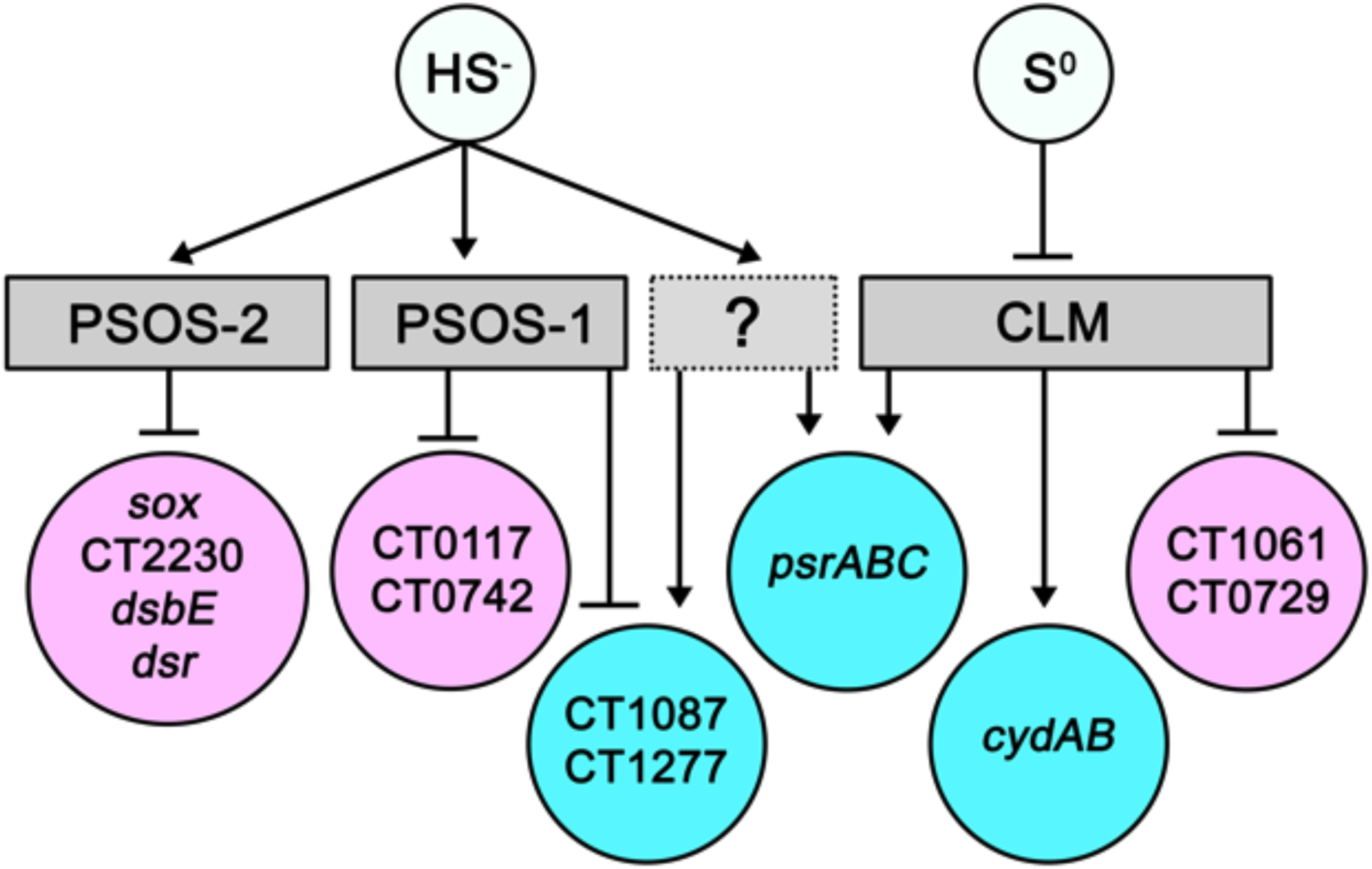
Simplified model of the oxidative sulfur metabolism regulatory network in *Cba. tepidum* with nodes representing metabolites (light blue), putative regulatory signals (gray), genes downregulated on sulfide relative to S(0) (pink), and genes upregulated on sulfide relative to S(0) (blue). Arrows represent activation, and bars represent repression.

For several of the key components of sulfide oxidation that are downregulated in response to sulfide, a putative operator sequence, PSOS-2, was discovered overlapping the TSS and/or the RpoD -6 box (Fig. 3). The positioning of this motif, and the expression patterns of the associated transcriptional units, strongly suggest that this motif functions as an operator sequence. As these genes are downregulated in response to sulfide relative to thiosulfate (5), and appear to also be downregulated in response to sulfide on S(0) (Fig. 3), PSOS-2 may bind a sulfide-dependent repressor (Fig. 6). CT2230 is predicted to be a membrane transporter in the FadL family of outer membrane proteins, and has been characterized in long chain fatty acid transport in *E. coli* (30). As CT2230 is involved in the transport of hydrophobic molecules, and is predicted to be part of a sulfide-repressed regulon, it may be involved in transport of hydrophobic sulfur chains across the outer membrane.

The *psrABC* promoter was found to contain two CLM sites, with single CLM sites occurring near the TSS of *cydAB*, CT1089-CT1088, CT1061, and CT0729 (Fig. 4). In *E. coli*, CRP can function as both a repressor or activator depending on positioning relative to the TSS and sigma factor binding site (21). Both *psrABC* and *cydAB* displayed similar expression profiles in that they were downregulated on S(0) relative to thiosulfate and sulfide (Fig. 4D). However, CT1089-CT1088 did not change significantly in expression between growth conditions, and CT1061 and CT0729 are both upregulated on S(0) relative to sulfide. These expression patterns can be explained if the location of the CLM site is taken into account (Fig. 4A,B), and if S(0) inhibits the activity of the CLM-binding protein. For *psrABC* and *cydAB*, the CLM sequence likely functions as an activator, while for CT1061, the oTSS, and CT0729 it likely functions as a repressor in that it interferes with sigma factor binding or transcription elongation. The CLM site may not have a large effect on CT1089-CT1088, and could function as either repressor, or activator (21). Thus, if S(0) inhibits the function of the CLM, it would repress *psrABC* and *cydAB*, while activating CT1061 and CT0729 (Fig. 6).

Between the transcript abundance data (this study; 5), and the three putative regulatory motifs discovered in sulfur-regulated genes, it appears that sulfide likely acts as a master regulator, activating proteins that bind PSOS-1, where CT1277 is the most likely candidate binding protein, PSOS-2, and the inferred, but unidentified, sulfide-dependent activator acting on CT1087 (Fig. 6). If S(0) inhibits the activity of the CLM binding protein, where CT1719 is the most likely candidate binding protein, then the expression pattern of CLM-associated genes can be explained. As the three motifs identified here were found to be conserved between multiple *Chlorobiaceae* genomes, it suggests that the regulatory functions carried out by these motifs are conserved across genomes. Thus, we have identified genes important for growth on S(0) relative to sulfide and thiosulfate, globally mapped TSS that were active during growth on these electron donors, identified basal promoter motifs, and discovered putative sulfur regulatory motifs. From this data, a model has emerged whereby sulfide is a master regulator of *Cba. tepidum*’s metabolic state, inducing the repression of genes important for thiosulfate and S(0) oxidation, and the activation of several key components for sulfide oxidation.

## Methods and Materials

### RNA sequencing

Cultures were grown at 20 µmol photon m^-2^ s^-1^ PAR in Pf-7 medium (24) at 47°C with thiosulfate as the sole electron donor, or at 42°C with biogenic S(0) as the sole electron donor. RNA was extracted using the NucleoSpin RNA kit (Machery-Nagel), rRNA was depleted using the MicrobExpress kit (Ambion), and treated with the TURBO DNA-free kit (Ambion) to remove residual genomic DNA. RNA-seq libraries were constructed using the NuGEN Ovation kit (NuGEN) to convert 25 ng RNA to double stranded cDNA, with subsequent fragmentation to approximately 100 bp fragments using an S2 Adaptive Focused Acoustic Disruptor (Covaris). Fragment size was confirmed by agarose gel electrophoresis on a 2% gel. The NuGEN Encore kit (NuGEN) was used to ligate adapters suitable for Illumina sequencing to the ends of these fragments. Libraries were sequenced on an Illumina HiSeq 1000 at the University of Delaware Sequencing and Genotyping Center.

### RNA-seq analysis

Reads were aligned to the *Cba. tepidum* genome using the Eland pipeline (Illumina). Custom Perl scripts (available upon request) were used to calculate the number of sequences that mapped to each annotated gene, non-coding RNA, and intergenic region of more than 50 bp, correcting for the length of the region. Data were normalized using quantile normalization as previously described (5). Differential expression between each pair of libraries was calculated using DESeq (30).

We previously reported the transcriptional response of *Cba. tepidum* to sulfide addition after growth on thiosulfate (5). The sulfide and S(0) expression libraries were constructed with different kits, and sequenced at different times, so in order to compare expression of *Cba. tepidum* on S(0) to that on sulfide, fold changes on S(0) relative to thiosulfate were divided by fold changes on sulfide relative to thiosulfate, giving a fold change value for each gene on sulfide relative to S(0). JMP® Pro (version 12.1.0) was used to create a box-and-whisker plot for the distribution of the fold change of sulfide relative S(0). Outliers, or those points outside of 1.5 × the interquartile range from the mean, were called as differentially expressed genes (Table S1).

### Differential RNA sequencing

dRNA-seq libraries were constructed from four independent cultures grown on thiosulfate, biogenic S(0), and from those grown on thiosulfate, spiked with 1.6 mM sulfide, and harvested 30 minutes after sulfide addition. *Cba. tepidum* cultures were grown as described by Levy, Lee, and Hanson at 20 µmol photon m^-2^ s^-1^ PAR (32). Replicate 1.5 ml cell pellets were harvested by centrifugation after 20 hours of growth (mid-log phase). Cell pellets were flash frozen in liquid nitrogen, and stored at -70°C. Cell pellets were thawed on ice, resuspended in 100 µl TE buffer (pH 8.0) with 1 µl Ready-Lyse Lysozyme (20 KU/µl; Epicentre), and incubated at room temperature for 30 min. Two replicate cell pellets were combined and RNA was purified by the NucleoSpin RNA kit (Macherey-Nagel). 10 µg RNA was treated with the TURBO DNA-free Kit (ThermoFisher) for gDNA removal, and then concentrated over RNA Clean & Concentrator-25 columns (Zymo Research). rRNA was depleted via the MicrobExpress kit (Ambion), and samples were concentrated over Zymo-25 columns. Libraries were normalized by *sigA* copy number (5) prior to terminator 5’ phosphate-dependent exonuclease (TEX; Epicentre) treatment. Each replicate was split into two samples for TEX treatment: the first was treated with 1 U TEX in a 20 µl reaction, while the second was incubated in the same buffer without TEX (33). Reactions were cleaned over RNA Clean & Concentrator-5 columns (Zymo Research). Samples were subsequently treated with 2 Units tobacco acid pyrophosphatase (TAP; Epicentre) in 50 µl total reaction volume followed by RNA concentration on Zymo-5 columns. 5’ adapters from the NEBNext Multiplex Small RNA Library Prep Set for Illumina kit (NEB) were ligated to the treated RNA samples using 2.5 µl Ligation Enzyme Mix in 20 µl total reaction volume with final concentrations of 1 mM ATP and 12.5% PEG8000. RNA was subsequently fragmented for 2 min using the NEBNext Magnesium RNA Fragmentation Module (NEB), and cleaned using Agencourt RNAClean XP beads (Beckman Coulter). 3’ ends were repaired via calf intestinal alkaline phosphatase (NEB) treatment, and cleaned over Zymo-5 columns. 3’ adapter ligation, first strand synthesis with indexed primers, and cDNA amplification (15 cycles) were completed using the NEBNext Small RNA kit. cDNA reactions were purified via Agencourt AMPure XP beads (Beckman Coulter). BluePippin was used for size selection (150-600 bp) prior to Illumina sequencing over two lanes (12 libraries pooled per lane) on a HiSeq 2000 at the University of Delaware Sequencing and Genotyping Center.

### dRNA-seq analysis

The RNA-seq analysis pipeline READemption (version 0.3.9) was used to align reads to the *Cba. tepidum* genome, and the output was passed to TSSpredator (version 1.06) for TSS identification (28, 33). In order for a TSS to be called as active under a specific condition, it needed to be detected in at least three of four replicates with an enrichment score of ≥2, and meet the remaining parameters specified under the *very specific* setting. TSS were classified according to the definitions of Sharma *et al.* (33). Primary TSS (pTSS) are defined as a TSS within 300 bp upstream of an annotated gene. If multiple TSS are present in this range, the pTSS is that with the highest enrichment value, and the others are secondary TSS (sTSS) sites. Antisense TSS (asTSS) occur on the antisense strand internal to or within 100 bp downstream of an annotated feature. Internal TSS (iTSS) occur on the sense strand within an annotated gene. Orphan TSS (oTSS) do not fall into any of the other class definitions.

### Motif discovery and analysis

Basal promoter motifs were identified by analyzing 50 bp upstream of all TSS by the MEME software suite (7). The output from multiple MEME runs from varying parameter settings were pooled via a custom R script (available upon request) to obtain estimates for the number of TSS associated with each motif. Promoter regions for orthologs of genes that are strongly regulated by sulfide in *Cba. tepidum* were extracted from all *Chlorobiaceae* genomes (Table S4). Up to 1000 bp upstream of the start codon for each orthologous gene set were analyzed by MEME and/or DMINDA (7, 35). Motifs identified as above were searched against the *Cba. tepidum* genome using FIMO to identify additional occurrences (15).

### Deletion mutagenesis

CT1277 was deleted from the *Cba. tepidum* genome using a counter-selectable suicide vector that will be fully described elsewhere (Hilzinger and Hanson, Unpublished). Briefly, a PCR product containing flanking DNA upstream and downstream of CT1277 was cloned into a mobilizable suicide vector with both antibiotic resistance and a counter-selectable marker based on a vector used for gene deletions in *Shewanella oneidensis* MR-1 (36). Primers used in this study are listed in Table S5.

### Response to sulfide qRT-PCR

*Cba. tepidum* wild-type and ΔCT1277 strains were grown in Pf-7 medium at 47°C and 20 µmol photon m^-2^ s^-1^ PAR. The absence of sulfide from cultures was verified by testing with CuCl_2_ (2). Pre-sulfide biomass was pelleted by centrifugation, flash frozen in liquid nitrogen, and stored at -70°C. Sulfide was added to a final concentration of 2 mM and cultures were incubated at 47°C and 20 µmol photon m^-2^ s^-1^ PAR for 40 min. Post-sulfide biomass was pelleted, flash frozen, and stored at -70°C. RNA was extracted, and purged of residual genomic DNA as described for dRNA-seq. *sigA* and CT1087 mRNAs were reverse transcribed into cDNA using the SigA-R-RT (24) and CT1087-R-RT primers (Table S5), and ProtoScript II cDNA synthesis kit (NEB) in the same reaction mixture. Negative controls lacking reverse transcriptase were performed to detect the presence of gDNA. The expression levels of *sigA* and CT1087 were determined using RealMasterMix SYBR ROX (5 Prime) on the ABI 7500 Fast Real-Time PCR System (Applied Biosystems). Genomic DNA standards were used to determine the efficiency of the SigA-RT and CT1087-RT primer sets and to quantify transcript abundance. CT1087 levels were normalized using *sigA* expression levels.

### Data deposition

Data described in this paper have been deposited in the National Center for Biotechnology Information Short Read Archive affiliated with BioSample accession numbers SAMN07413950 for S(0) and thiosulfate RNA-seq data and SAMN07413841 for all dRNA-seq data.

## Acknowledgements

This work was supported by NSF grant MCB-1244373 to T.E.H. and an IGERT SBE2 fellowship to J.M.H. This project utilized computational resources at the University of Delaware Center for Bioinformatics and Computational Biology Core Facility funded by Delaware INBRE (NIGMS GM103446), Delaware EPSCoR (NSF EPS-0814251, NSF IIA-1330446), the State of Delaware, and the Delaware Biotechnology Institute. The authors would like to thank Katie Kalis and Amalie Levy for helpful discussions during data analysis and manuscript preparation, and the University of Delaware Sequencing and Genotyping Center staff for advice and help with library construction and sequencing.

## Legends for Supplementary Material

**Table S1.** Differential gene expression analysis for RNA-seq data in Excel format (.xlsx). Sheet 1 (S0 vs. TS) contains DESeq output comparing RNA-seq libraries prepared from cells grown on S(0) (baseMeanA) vs. thiosulfate (baseMeanB) as the electron donor. Sheet 2 contains the analysis of expression ratios on different electron donors vs. thiosulfate (i.e. sulfide vs. thiosulfate compared to S(0) vs. thiosulfate) to identify genes differentially expression between S(0) and sulfide. See Materials and Methods for details of calculations.

**Table S2.** Summary alignment statistics for each dRNA-seq library produced in this study.

**Table S3.** All TSS identified in this study and their classification.

**Table S4.** *Chlorobiaceae* genome annotations used in the phylogenetic footprinting analysis.

**Table S5.** Sequences for oligonucleotide primers used in this study.

